# Inference of an integrative, executable network for Rheumatoid Arthritis combining data-driven machine learning approaches and a state-of-the-art mechanistic disease map

**DOI:** 10.1101/2021.01.28.428679

**Authors:** Quentin Miagoux, Vidisha Singh, Dereck de Mézquita, Valerie Chaudru, Mohamed Elati, Elisabeth Petit-Teixeira, Anna Niarakis

## Abstract

Rheumatoid arthritis (RA) is a multifactorial, complex autoimmune disease that involves various genetic, environmental and epigenetic factors. Systems biology approaches provide the means to study complex diseases through the integration of different layers of biological information. Combining multiple data types can help compensate for missing or conflicting information and limit the possibility of false positives. In this work, we aim to unravel mechanisms governing the regulation of key transcription factors in RA and derive patient-specific models to gain more insights into the disease heterogeneity and the response to treatment.

We first make use of publicly available transcriptomic datasets (peripheral blood) relative to RA and machine learning to create an RA specific transcription factor (TF) co-regulatory network. The TF cooperativity network is subsequently enriched in signalling cascades and upstream regulators using a state-of-the-art, RA-specific molecular map. The integrative network is then used as a template to analyze patients’ data regarding their response to anti-TNF treatment and identify master regulators and upstream cascades affected by the treatment. We use the Boolean formalism to simulate in silico subparts of the integrated network and identify combinations and conditions that can switch on or off the identified TFs, mimicking the effects of single and combined perturbations.

## Introduction

Rheumatoid Arthritis (RA) is an inflammatory, autoimmune disease that affects the joints of the body. While the exact aetiology is unknown, it involves a combination of environmental and genetic factors such as smoking and the presence of susceptibility genes, along with sex and age factors. RA affects 0.5-1% of the world population with women three times more susceptible to develop RA than men [1,2]. The onset of the disease is set around the fourth to fifth decade of one’s life [3] and if left untreated can be debilitating for the individual.

Symptoms of RA include synovial inflammation, joint stiffness and pain, cartilage destruction and bone erosion. In early RA, leukocytes invade the synovial joints followed by other pro-inflammatory mediators which instigate an inflammatory cascade and provoke synovitis [2]. Activated monocytes and T cells, both a source of proinflammatory cytokines such as TNF-a, can be found in peripheral blood [4] and many RA studies have used peripheral blood cells to identify disease-related genes [5– 8]. The common therapy for the RA includes the use of Disease-modifying anti- rheumatic drugs (DMARDs). Conventional DMARDs include drugs that target the entire immune system whereas biologic DMARDs are monoclonal antibodies (mAbs) and soluble receptors that target protein messenger molecules or cells. Patients who do not respond to conventional DMARDs usually initiate therapy with TNF inhibitors. However, approximately 30–40% of RA patients fail to respond to antiTNF therapy and are usually obliged to undergo several rounds of drug combinations [9].

Due to the complex nature of RA, systems biology and integrative approaches are needed in order to gain insight into the disease pathogenesis and progression. It is evident that focusing only on one aspect of the disease provides a limited understanding of the multifactorial nature of RA. Recently, many computational approaches, mainly network-based which rely on the integration of multi-omics data (proteomics, genomics, transcriptomics and metabolomics) have succeeded in unravelling key mechanisms in complex diseases [10–13]. In this direction, machine learning is a promising bioinformatics field that allows the use and integration of a variety of biomedical data with inherent complexity and large size. Studies have shown that incorporating prior knowledge to data-driven methodologies improves the quality and the biological relevance of the outcome [14–16]. One such machine learning tool is CoRegNet, an R/Bioconductor package that infers co-regulatory networks of transcription factors (TFs) and target genes by analyzing transcriptomic data and estimating TFs activity profiles. Moreover, the software also allows for network enrichment by integrating regulation evidence for TF binding sites, protein-protein interaction data and ChiP data from various databases to support cooperative TFs [17].

In this work, we present a framework for the integration of signalling and transcriptional regulation cascades with genomic mutations, combining data-driven approaches with prior knowledge in the form of an integrative RA-specific network. To do so, we make use of publicly available transcriptomic data of white blood cells from patients suffering from RA, and the tool CoRegNet to infer a co-regulatory network. Next, we develop an integration pairing method to couple the RA co-regulatory network with a state-of-the-art disease map for RA [18], to enrich the cooperativity network with upstream, signalling regulators. Disease maps are comprehensive, knowledge-based representations of disease mechanisms including disease-related molecular interactions supported by literature-based evidence [19,20].

Next, we project on the integrative RA network public genomic data and transcriptomic data from treated RA patients, highlighting key mutation carriers and differentially expressed genes associated with the response to anti TNF treatment (Figure 1). The goal is to unravel mechanisms governing the regulation of key transcription factors and genes identified as mutation carriers or DEGs in RA patients undergoing anti TNF treatment. Lastly, we study the dynamic behaviour of the system using the Boolean formalism to simulate subparts of the integrated network [21,22]. This includes real time simulations, sensitivity analysis and dose-response analyses to study the impact of other signaling cascades on the expression of the identified TFs, and steady state analysis revealing combinations and conditions that can switch on or off the identified TFs, mimicking the effects of the treatment [23].

**Figure 1.**
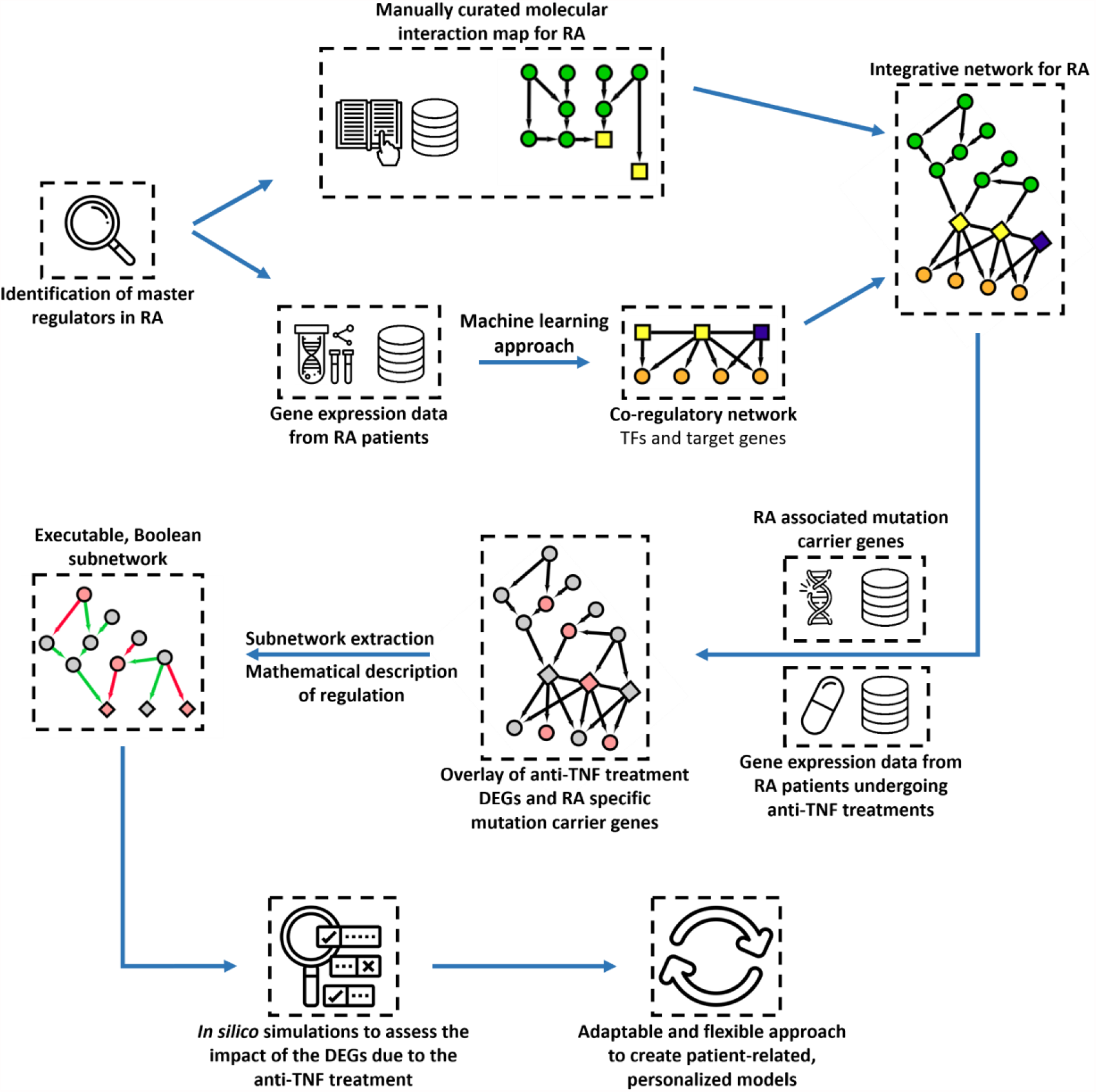
Workflow of an integrative and executable network for Rheumatoid Arthritis. From the identification of master regulators in Rheumatoid Arthritis to an adaptable and flexible personalized model, combining data-driven machine learning approaches and a state-of-the-art mechanistic disease map.

## Materials and Methods

### Data description and pre-processing

We used a transcriptomic dataset from Illumina HiSeq 2500 of white blood cells from RA patients and healthy donors (GSE117769). The data set describes 120 subjects in total (51 RA and 50 controls, and 19 patients with either Ankylosing spondylitis or Psoriatic arthritis). From this dataset, we extracted 96 samples (46 RA and 50 controls). For the 46 RA samples, 43 were of Caucasian ancestry and 3 of unknown ancestry. Moreover, two samples consisted of duplicates of the same origin (female, control, unknown ancestry) and their expression matrix average was used for the analysis. We conducted a preliminary analysis on the expression matrix data with DESeq2 [24], using normalization and variance stabilizing transformation on the matrix expression. Principal Component Analysis (PCA) revealed 5 outliers (4 RA and 1 control) that were removed from the dataset (Figure S1). The final dataset used for further analysis comprised 90 samples (42 RA and 48 controls). Finally, we performed a normalisation on the raw expression matrix after removal of low read counts (>10).

### Inference of the co-regulatory network (CoRegNet)

We used CoRegNet [17] R package version 1.26.0 to infer the co-regulatory network with the normalised gene expression matrix of the pre-processing step. The CoRegNet package implements the H-LICORN algorithm, allowing the identification of cooperative regulators of genes [25]. We enriched the inferred network with protein-protein and regulation evidence and further refined it with an unsupervised method using the unweighted mean. The co-regulatory network inferred with CoRegNet is composed of the significant edges between TF with a False Discovery Rate (FDR) [26] of 5%.

### RA map upstream protein extraction

The RA map is a state-of-the-art interactive knowledge base for the RA disease [18]. The RA map is organized in the form of a cell representing the flow of information from the extracellular space to the plasma membrane and then to the cytoplasm, the nucleus and the secreted compartment or cellular phenotypes. For our analysis, as we were focused on upstream regulators of identified TFs, we used mainly the signalling part of the RA map.

More specifically, the list of TFs was uploaded as an overlay to the RA map, and the matching TFs were identified. The matching TFs were subsequently used as seeding nodes for the upstream plugin [27], setting the mode of extraction as upstream and selecting non-blocking modifiers. The obtained file was extracted as an XMLCellDesigner file. With this method we extracted upstream signalling cascades, and also seven translation reactions for which the mRNA was directly linked with the protein in the cytoplasm or the membrane.

The RA map, and consequently the extracted network, is written in the Process Description Systems Biology Graphical Notation scheme [28]. To obtain a more simplified, Activity Flow (AF) - like representation of the network, the CellDesigner XML file of the previous step was used as an input to the tool CaSQ [29] to create an Activity Flow like executable network. CaSQ provides SBML-qual files for performing *in silico* simulations, but in our case, we used only the SIF file that contains information about the source, interaction type, and target of the Boolean network.

The obtained SIF file was further modified to address the issue of complexes - for every complex represented as a single node in the AF network, we recreated the reactants. This way we would not miss interactions and overlaps between nodes existing inside complexes. Regarding entities represented multiple times (as genes, proteins or mRNAs), for simplification purposes we kept only one entity and merged the corresponding interactions.

### Global RA Network inference

We used the R package igraph [30] to convert the CoRegNet object (co-regulatory network) and the RA map SIF file into separate graphs. Then, we merged both networks using igraph, and imported the network into Cytoscape using the RCy3 R/ Bioconductor package [31] forming the global RA-specific network.

### Differential expression analysis (DEA) using independent datasets

We conducted multiple differential expression analyses (DEA) using two different datasets. One dataset contained normalised counts from RNA sequencing data of CD4+ T cells including different responses to anti-TNF on RA patients (GSE138747). 37 and 41 RA patients treated with Adalimumab and Etanercept compose this dataset respectively, which were analysed independently. The second is composed of raw counts from RNA sequencing data of whole blood cells of biologic naive rheumatoid arthritis patients from baseline and after 3 months treatment with Infliximab or Adalimumab (GSE129705). This dataset contains two different cohorts of 40 and 36 RA patients, which were also analysed independently. Thus, using DESeq2, we conducted two DEA on the comparison of responders and non-responders gene expression level for both drugs (Adalimumab and Etanercept) and two DEA on the comparison of baseline and after 3 months of anti-TNF treatment gene expression level. Therefore, from all these analyses, we considered a gene as a differentially expressed gene (DEG) with a corrected p-value (FDR) < 0.1. Finally, DEGs from both analyses were used as an overlay for the global RA-specific network.

### List of variants

DisGeNET [32] contains the largest publicly available collection of genes and variants associated with human diseases. From this database, we extracted 2387 variants associated with RA. Then, we filtered out variants that had a variant disease association (VDA) and evidence index (EI) score lower than 0.7. VDA score is computed using the number of curated and non-curated publications supporting the variant disease association, while EI score is computed using contradictory results in publications supporting the variant. The use of a 0.7 threshold gives us at least one curated publication supporting the variant and the disease association resulting in a total of 1635 variants. Within these 1635 variants, we identified 731 associated genes that were subsequently used as an overlay for the global RA-specific network.

### Subnetwork extraction

The subnetwork, based on the global network for RA, is focused on TNF, IL6 and TGFB1, 3 molecules highly implicated in RA (see Discussion). These 3 proteins were extracted along with their downstream cascades up to the first TF, to reduce complexity and focus on the upstream regulators. From the global network for RA displayed in Cytoscape, we selected TNF, IL6 and TGFB1 simultaneously and using the Biological Network Manager (BiNoM) plugin, we selected in a stepwise manner the downstream neighbors of TGFB1, IL6 and TNF up to the first affected TF(s).

### Shiny app

The co-regulatory network inferred with CoRegNet, the RA map Activity Flow extracted network and the merged global RA network, along with their overlays, were integrated to a web-based Shiny application using R [33]. The web application uses a Cytoscape viewer based on the R package cyjShiny [34] and is freely available (https://quentin-miagoux.shinyapps.io/global_ra_network/).

### Subnetwork logic-based dynamical

The subnetwork previously obtained from the global network for RA, displayed on Cytoscape was extracted in SBML format using the BiNoM plugin and its function “export to SBML”. Then, we removed co-regulatory interactions between TFs, which could have been misleading and interpreted as activation in further boolean simulation. We adjusted the overlay using CellDesigner and converted the SBML file into SMBL- Qual using CaSQ (v0.9.11).

The SMBL-Qual file was then imported on CellCollective to perform real time simulation. Using the “Simulation” tab on CellCollective, we mimicked the downregulation according to each subnetwork conditions (before/after treatment, responders/non-responders, mutation carriers), and activated one by one each input (TNF, IL6 and TGFB1). Furthermore, we also performed a sensitivity and dose response analysis with CellCollective, using 5 different initial states, described in Table 3.

The same file was used for analysis with the software GINsim [35] after a post processing modification step for node name recognition by the software. We used the nightly build version 3.0.0b-SNAPSHOT and the functions *Reduce model* for the reduced version, *Compute stable states* to obtain the stable states of the model and *Run simulation* with configurations of perturbations for the *in silico* KO experiments.

## Results

### Inference of the co-regulatory network

To infer the co-regulatory network, we selected a transcriptomic dataset from GEO database (GSE117769), including 120 samples (51 RA and 50 control, and 19 patients with either Ankylosing spondylitis or Psoriatic arthritis). After a series of pre-processing checks including the sample origins, duplicates, and quality of the data using a PCA on the matrix expression with a normalization and variance stabilizing transformation (shown in Figure S1), we ended up with 90 patients (48 Controls and 42 RA patients). Then, of the remaining samples we obtained normalized counts using DESeq2, on which we finally applied CoRegNet to infer the co-regulatory network, which is presented in Figure 2.

**Figure 2.**
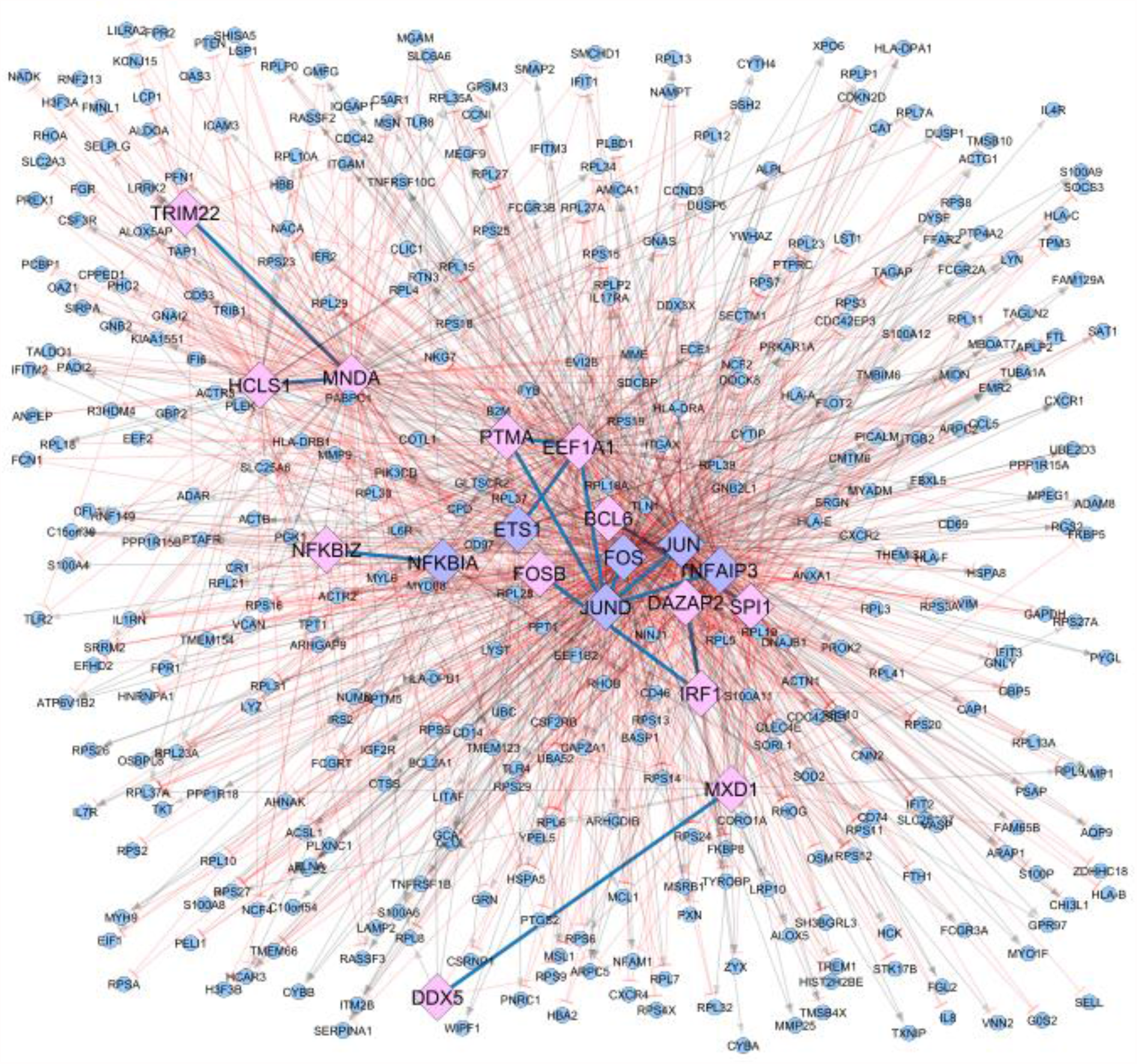
Co-regulatory network inferred using the tool CoRegNet and the matrix of normalized counts from the transcriptomic dataset (GSE117769). The dataset used included data from 90 patients (48 Controls and 42 RA patients). Matching TF from CoRegNet with the RA map are coloured in purple (6) while non-matching are colored in pink (13). CoRegNet target genes are coloured in blue (373). Inhibitions are represented with red blunt arrows, and activations with grey arrows.

This network includes a total of 19 TF, 14 co-regulatory interactions and a total of 373 regulated target genes. Table 1 summarizes the top five TFs with the highest number of regulatory and co-regulatory interactions. Literature search for the nineteen TF identified from CoRegNet as the master regulators in the dataset, showed their potential implication to RA. Supplementary Table S1 summarises key roles of the TFs and the corresponding literature reference.

**Table 1.** Top 5 of the CoRegNet identified TFs with the highest number of regulatory interactions (TF - target gene) and co-regulatory interactions (TF-TF).

| **Top 5 TFs** | **No of regulatory interactions** |
| --- | --- |
| FOS | 288 |
| JUN | 211 |
| EEF1A1 | 155 |
| MNDA | 136 |
| TNFAIP3 | 125 |
| **Top 5 TFs** | **No of co-regulatory interaction(s)** |
| JUND | 5 (EEF1A1, FOS, JUN, PTMA, TNFAIP3) |
| EEF1A1 | 3 (ETS1, JUND, PTMA) |
| IRF1 | 2 (DAZAP2, FOSB) |
| MNDA | 2 (HCLS1, TRIM22) |
| PTMA | 2 (EEF1A1, JUND) |

### RA map upstream regulators of the TFs identified from CoRegNet

Six out of the 19 TFs namely ETS1, FOS, JUN, JUND, NFKBIA and TNFAIP3 from the co-regulatory network were present in the RA map and were used as seeds to extract their upstream regulators. The extracted network comprising the RA map upstream regulators of the matching TFs includes 244 nodes and is shown in Figure 3.

**Figure 3.**
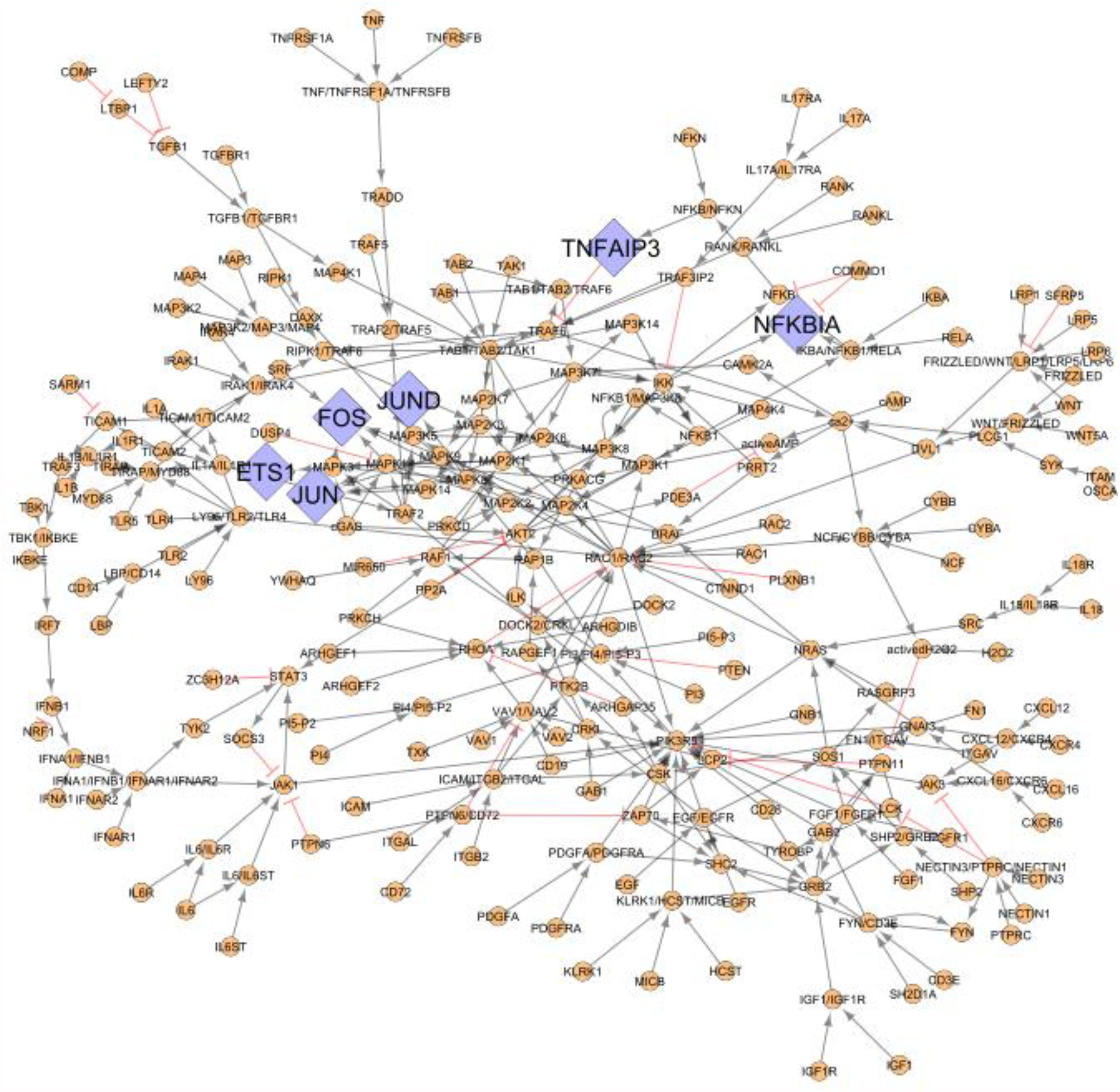
The RA map upstream regulators of the matching TFs between the RA map and the CoRegNet co-regulatory network. Matching TFs from the inferred network with CoRegNet are coloured in purple (6) and upstream regulators are coloured in orange (238). Inhibitions are represented with red blunt arrows, and activations with grey arrows.

### Coupling gene co regulation with signalling cascades to obtain a global, integrative RA network

The global, integrative RA network is the result of the merging of the RA map signalling cascades and the CoRegNet object, using as interface the matching TFs. It comprises 614 nodes and 1736 interactions (848 inhibitions, 874 activations, and 14 co-regulatory interactions shared among TFs) including genes, proteins, complexes, and simple molecules shown in Figure 4. In this network, six TFs were common between the CoRegNet network and the RA map (seeding TFs). In addition, 16 target genes identified with CoRegNet overlapped with the RA map upstream regulators.

**Figure 4.**
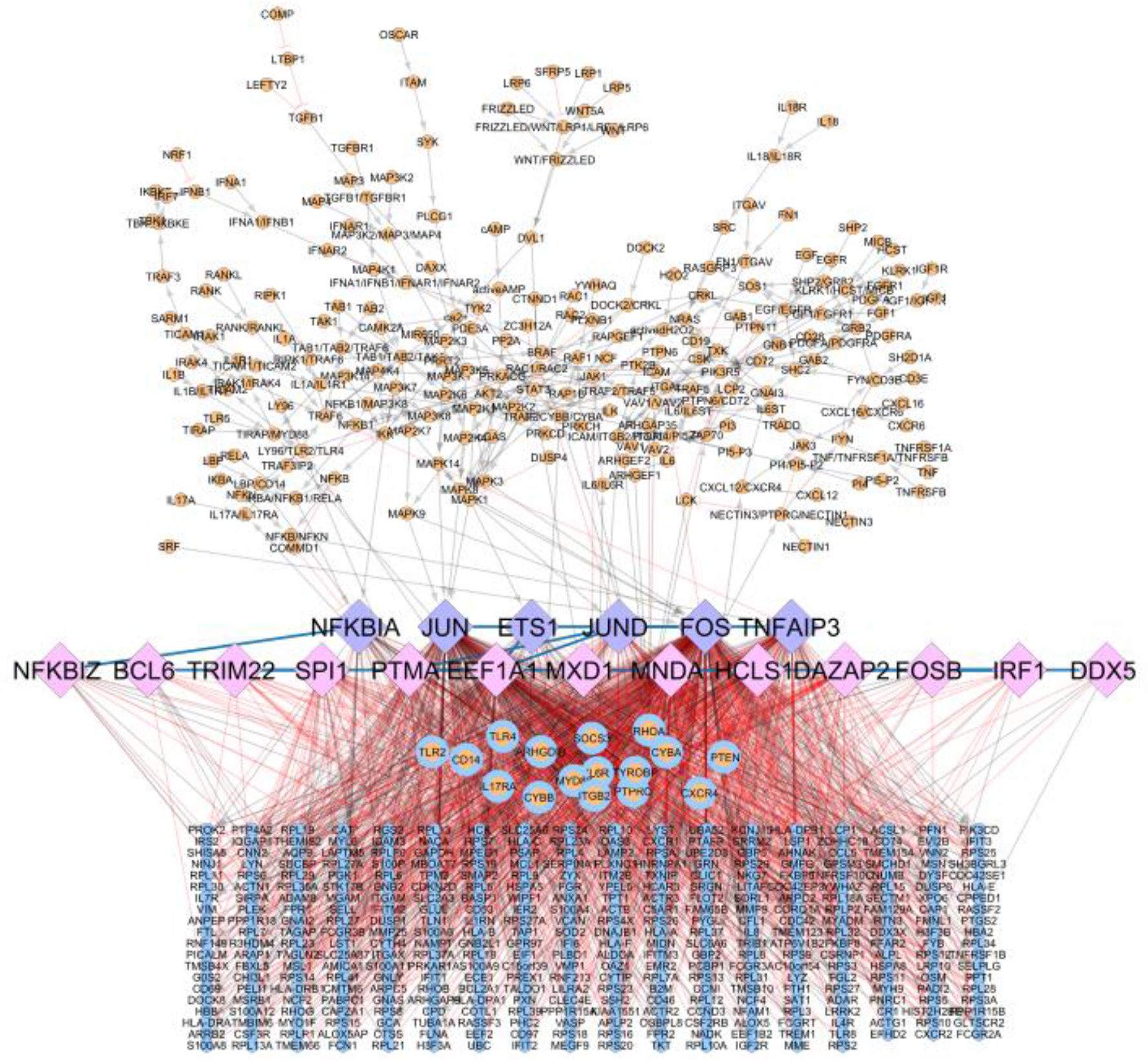
Integrative global network for RA. RA map upstream regulators are bound to TFs identified in the CoRegNet co-regulatory network. Matching TFs are coloured in purple (6), matching genes/proteins are coloured in blue and orange (16), upstream regulators from RA map are coloured in orange (222), CoRegNet TFs are coloured in pink (13) and CoRegNet target genes are coloured in blue (357). Inhibitions are represented with red blunt arrows, and activations with grey arrows.

### Two use-cases: Identification of key TFs using DEG from RA patients undergoing anti-TNF treatment

Two datasets of RNAseq expression data coming from RA patients undergoing anti-TNF treatment were analysed to obtain differentially expressed genes (DEG). The first dataset focuses on responders and non-responders to RA treatment, including 37 and 41 RA patients treated with Adalimumab and Etanercept respectively. The second one involves untreated and treated (Infliximab or Adalimumab) RA patients including two cohorts of 40 and 36 RA patients, respectively. DEGs from these analyses were mapped to the global network for RA (presented in Figure S2 and Figure S3 respectively).

DEGs from responders/non-responders RA patients data mapping shows a total of 15 matching nodes, including 4 Etanercept DEGs and 11 Adalimumab DEGs. Also, 4 matching nodes are CoRegNet and RA map TFs (NFKBIA, JUN, FOS and TNFAIP3) and 1 CoRegNet TF only (FOSB), in the global network for RA. DEGs from untreated and treated RA patients mapping show a total of 101 matching nodes, including 2 CoRegNet and RA map TFs (NFKBIA and FOS) and 4 CoRegNet TF only (BCL6, MXD1, MNDA and DAZAP2).

Finally, cross-analysis revealed that a total of 9 over 19 TFs included in the global network for RA overlapped with a DEG from at least one analysis (presented in Table 2). Among these 9 TFs, two of them, NFKBIA and FOS overlapped with a DEG in both analyses.

**Table 2.** Transcription factors from the global network for RA with at least one differentially expressed gene (DEG) overlapping from the analysis of a) responders/non responders RA patients to anti-TNF treatment and b) before/after anti-TNF treatment of RA patients. ↓ denotes downregulation.

|  | TF | Responders /<br>non-responders | After treatment /<br>before treatment |
| --- | --- | --- | --- |
| **CoRegNet and RA map**<br>**TF** | *NFKBIA* | ↓ | ↓ |
|  | *JUN* |  | ↓ |
|  | *FOS* | ↓ | ↓ |
|  | *TNFAIP3* |  | ↓ |
| **CoRegNet TF** | *BCL6* | ↓ |  |
|  | *MXD1* | ↓ |  |
|  | *MNDA* | ↓ |  |
|  | *DAZAP2* | ↓ |  |
|  | *FOSB* |  | ↓ |

### Subnetwork logic-based dynamical analysis

While the analyses highlight TFs differentially expressed (downregulated) after treatment or response to treatment with anti-TNF drugs, it is evident from the global network that the identified TFs can be regulated by a variety of other upstream cascades, besides those implicated in the TNF signalling.

To study further the interconnections with other pathways, we focused on IL6 and TGF-beta signalling (mentioned as TGFB1 in the network). Interleukin-6 (IL6) is the target of Tocilizumab (TCZ), an IL6 inhibitor frequently used in the treatment of RA. The inhibitor was developed in 2008, and its efficacy in treating the disease is quite similar to those of tumor necrosis factor (TNF) inhibitors [36]. TGF-beta signalling has been shown to be activated in RA synovium, however TGF beta blockage did not seem to have an effect on experimental arthritis [37]. To study further the impact of these cascades on the expression of the identified TFs, we constructed a subnetwork, by selecting the molecules TGFB1, IL6 and TNF in the global network along with their downstream neighbors until reaching an identified TF.

The subnetwork contains 38 nodes, including 4 TFs and is highly enriched in MAPKs, as seen in Figure 5. By projecting the DEGs and known genomic variants associated with the disease we can see that intermediate nodes and TFs are downregulated by the anti-TNF treatment, while the inputs (IL6, TNF and TGFB1) along with a few intermediate nodes are characterized as mutation carriers.

**Figure 5.**
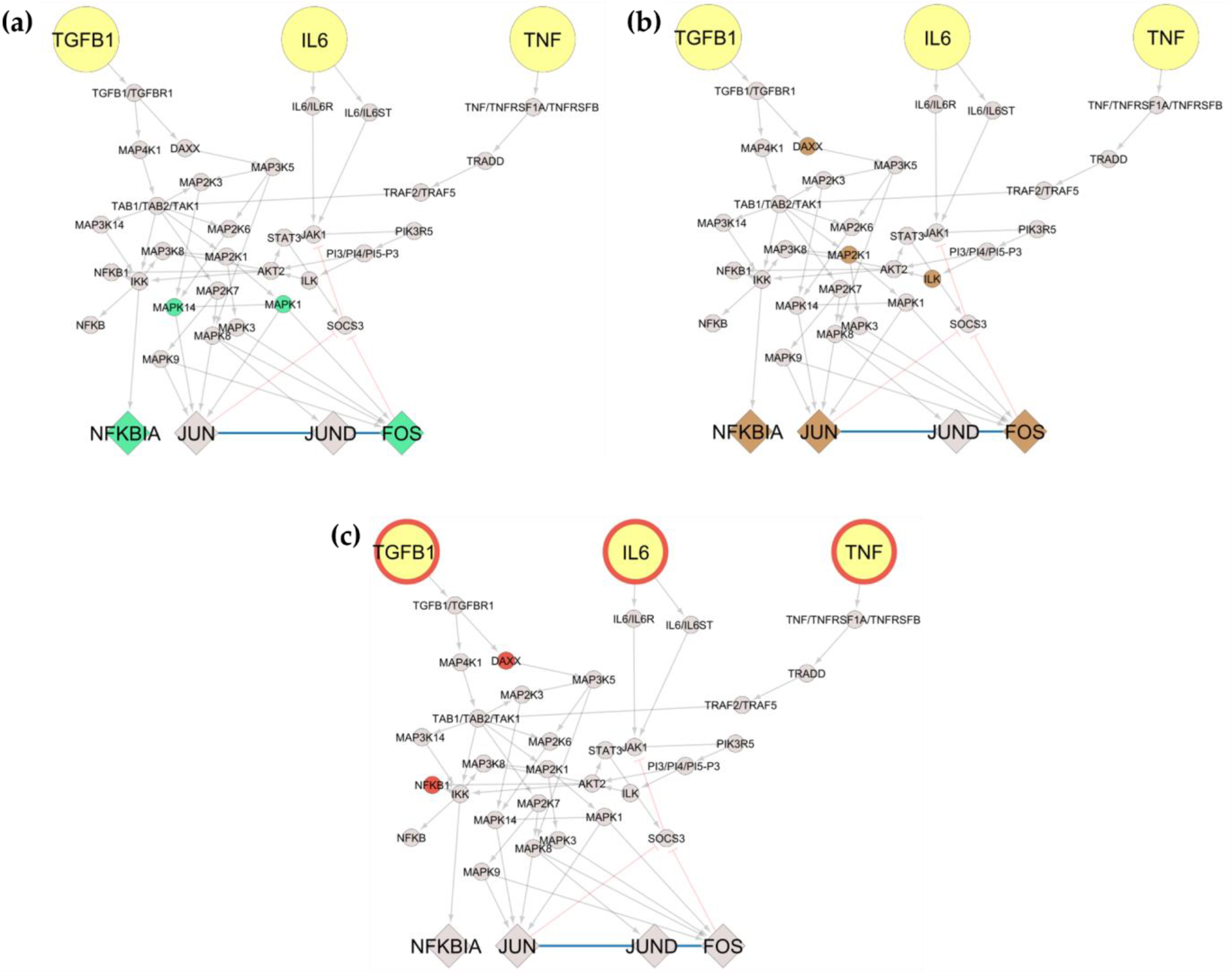
Subnetwork extraction of target genes (TGFB1, IL6 and TNF) from the global network for RA including differentially expressed genes and genomic variant overlap. We extracted three target genes (TGFB1, IL6 and TNF) with their downstream neighbors until the first transcription factor matches. Then, we overlapped over the subnetwork three analyses including **(a)** Matching differentially expressed genes (DEG) from responders/non responders RA patients to anti-TNF treatment shown in green (4). **(b)** Matching DEG before/after anti-TNF treatment of RA patients shown in brown (6). **(c)** Matching DisGeNET variants shown in red (5).

In the next step, we wanted to evaluate the impact of single and combined perturbations of the network inputs, on the expression of the TFs. To do so, we used the possibility of adding Boolean rules to the network with the tool CaSQ [29]. Boolean models have been long used to describe biological mechanisms in health and disease [38], and they are an optimal approach for modelling signalling and gene regulation, when kinetic parameters are scarce. Boolean models are qualitative by nature, based on the assignment of binary values to the variables and the use of the logical operators AND, OR and NOT to describe the regulation of all molecules in the system [21]. The CaSQ tool receives as an input an SBML CellDesigner file [39] and produces a Boolean network with preliminary logical rules.

The Boolean model produced from our subnetwork has 38 nodes (three inputs, six outputs and 29 intermediate nodes) and 59 interactions. The SBML qual file was imported to Cell Collective [40] to perform real time simulations and sensitivity analysis and to GINsim [35] to calculate stable states and perform *in silico* KO (knock out) simulations. For the *in silico* KO simulations, a reduced version of 23 nodes was used.

#### Real time simulations using the Cell Collective platform

First we wanted to see the impact of the molecules either affected by the treatment or identified as mutation carriers, to the model outputs. The analysis corresponding to before and after anti-TNF treatment, showed that MAPK14 and MAPK1 were downregulated. Accordingly, for the responders and non responders analysis, MAP2K1, ILK, DAXX were also identified as downregulated. Lastly, DAXX and NFKB1 were identified as mutation carriers. To mimic the effects of the downregulation of these molecules on the model outputs, we performed *in silico* simulations setting their activation level to zero.

For the dataset of before and after anti-TNF treatment of RA patients, MAPK14 and MAPK1 activity levels were set to zero, and simulations turning the inputs sequentially active revealed that when setting either TNF, TGFB1 or Il6 on, all TFs are expressed (Figure 6 a-c). When mimicking the downregulation of MAP2K1, ILK, DAXX for the dataset of responders/non responders to anti-TNF treatment, we observed that when setting TNF and TGFB1 on, all TFs are expressed (Figure 6 d,e). However, when IL6 is set on, only the TF NFKBIA is expressed (Figure 6 f). Finally, when mimicking the downregulation of DAXX and NFKB1 for the mutation carrier from DisGeNET, we observe that when setting either TNF, TGFB1 or IL6 on, all TFs are expressed (Figure 6 g-i).

**Figure 6.**
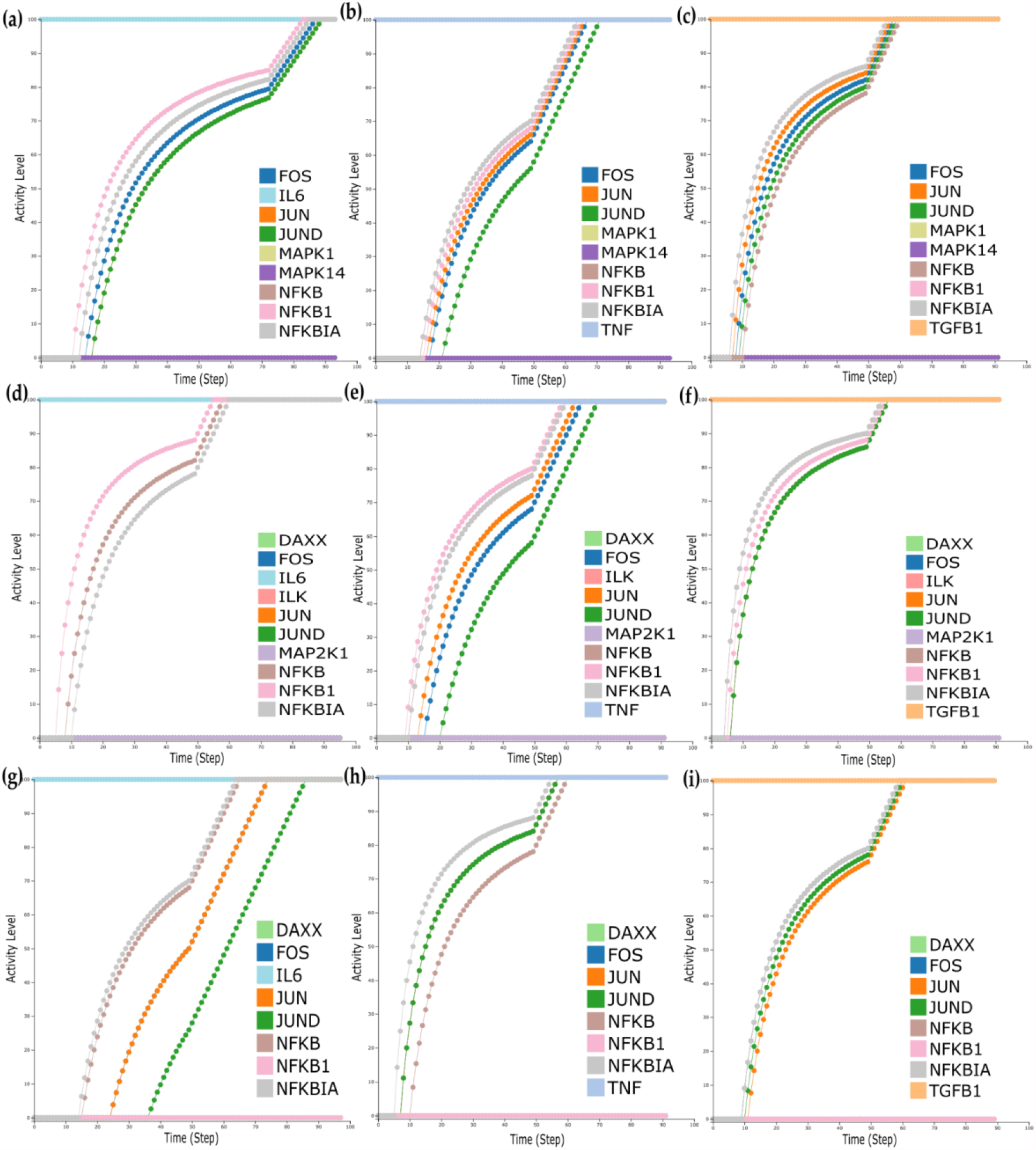
Real time simulations with Cell Collective. **1)** For the dataset of before and after anti-TNF treatment of RA patients, where MAPK14 and MAPK1 were found downregulated **(a-c). (a)** Simulations with IL6 activity set to 100%. **(b)** Simulations with TNF activity set to 100% **(c)** Simulations with TGFB1 activity set to 100%. **2)** For the dataset of responders/non responders where MAP2K1, ILK, and DAXX were found downregulated **(d-f). (d)** Simulations with IL6 activity set to 100%. **(e)** Simulations with TNF activity set to 100%. **(f)** Simulations with TGFB1 activity set to 100%. 3) For the dataset of the mutation carrier. **(g-i). (g)** Simulations with IL6 activity set to 100%. **(h)** Simulations with TNF activity set to 100%. **(i)** Simulations with TGFB1 activity set to 100%.

#### Dose response and Sensitivity analysis

For the dose response we studied five different initial conditions shown in Table 3 that mimic the effect of different scenarios in combination to TNF activity status. The first condition corresponds to TNF blockage and simultaneous impairment of IL6 and TGFB1 signalling, the second corresponds to having the IL6 cascade active, the third to having the TGFB1 active, the fourth both IL6 and TGFB1 active and lastly, the fifth condition mimics what happens to the system when only the TNF input is active.

We performed dose response analysis for all aforementioned conditions and observed that the expression of the TFs is dose dependent for TNF, TGFB1, and IL6 (Figure 7 b, e and f), while simultaneous activation of IL6 and TGFB1 cascades has a synergistic effect causing an increase of the activation levels of the TFs even for lower doses of IL6 and TGFB1 (Figure 7 c and d).

**Table 3.** Initial conditions for dose response analysis.

| **Initial conditions** | **Input state** |
| --- | --- |
| 1 | All inputs inactive |
| 2 | IL6 active |
| 3 | TGFB1 active |
| 4 | IL6+TGFB1 active |
| 5 | TNF active |

**Figure 7.**
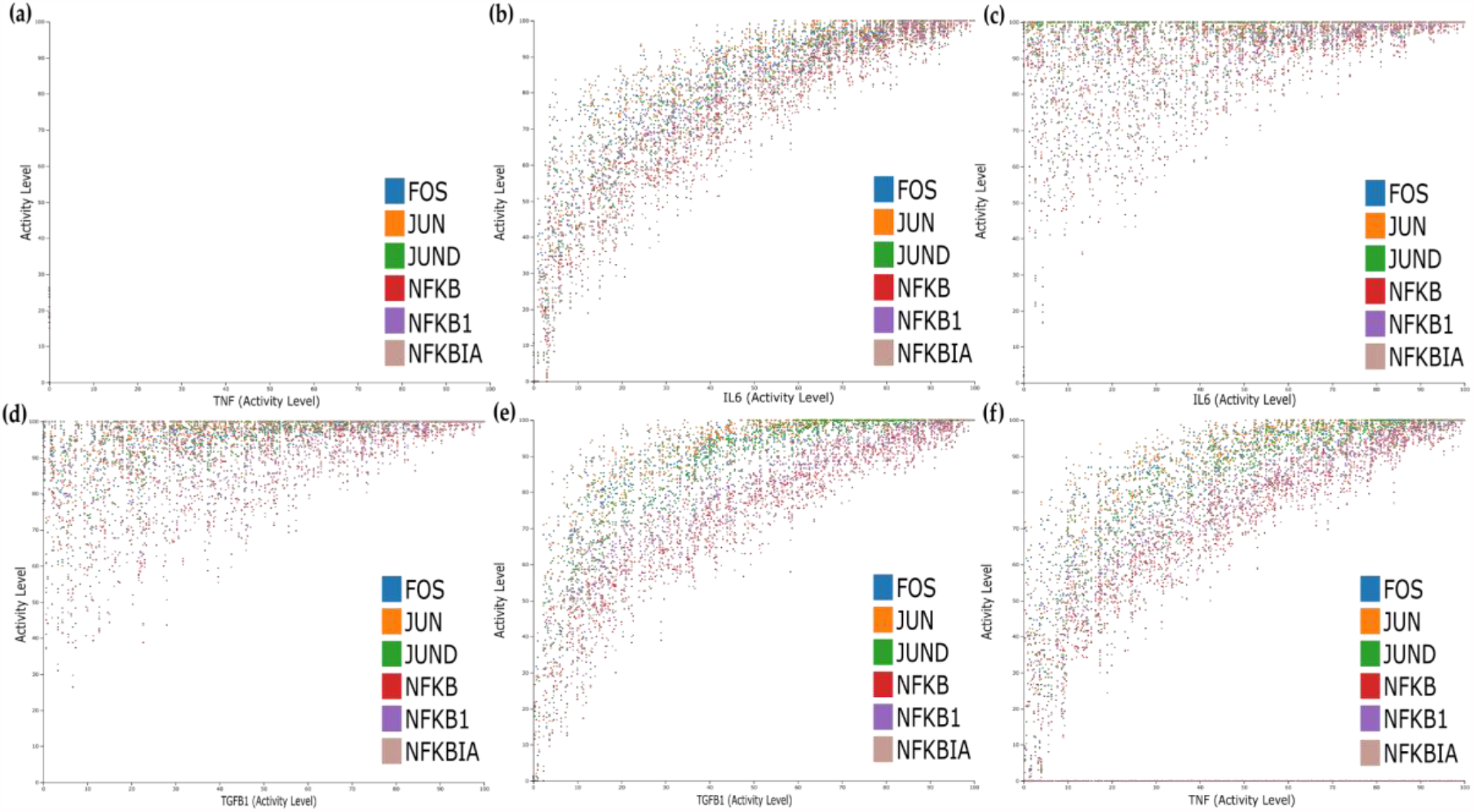
Dose response analysis. **(a)** All input inactives. **(b)** IL6 active. (**c)** IL6 and TGFB1 active (TGFB1 view). **(d)** IL6 and TGFB1 active (IL6 view). **(e)** TGFB1 active. **(f)** TNF active.

Next, we wanted to see how the downregulation of the TFs observed after anti TNF treatment could be counterbalanced by the other pathways, given that in non responders the expression of these TFs was kept intact, despite the treatment. We performed environmental sensitivity analysis to identify which of the two model inputs (IL6 and TGFB1) has the greatest impact on the up regulation of the TFs included in the model, when TNF activity is blocked. The results depicted in Supplementary files, showed that indeed the TFs could be upregulated in the absence of TNF activity, for a combination of activity ranges of the other two inputs (Suppl Figures S4).

#### Wild type stable state analysis and KO simulations

We performed stable state analysis for the model using the software GINsim. The analysis for the wild type (no perturbations) revealed five steady states (fixed points) and no complex attractor. The configurations of these five stable states as far as the molecules of interest are concerned (grey, blue and pink nodes of Figure 8) are shown in Table 4.

**Table 4.**
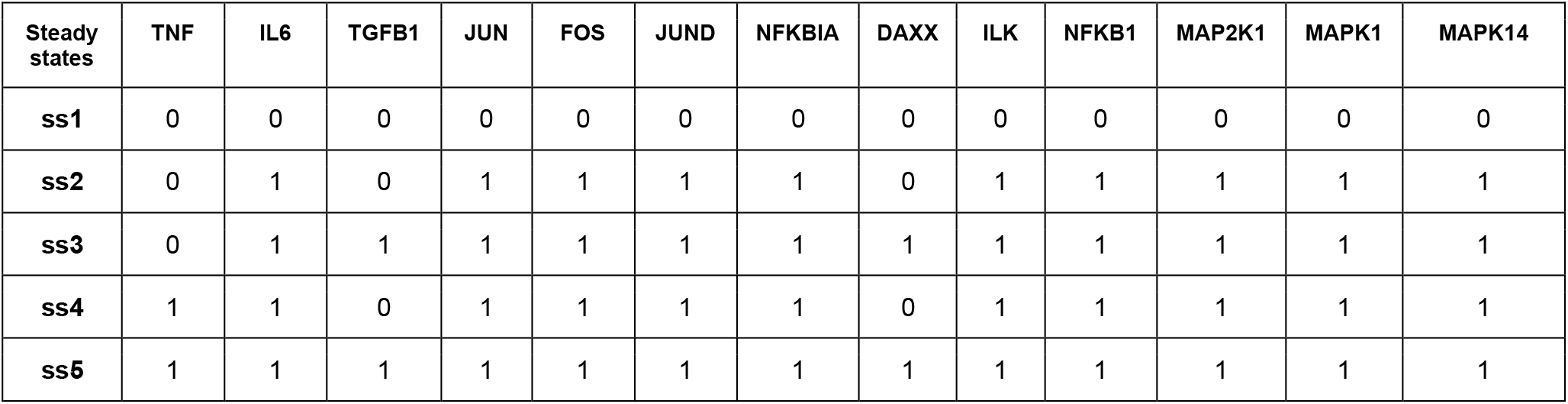
Stable states of the Boolean network (wild type).

**Figure 8.**
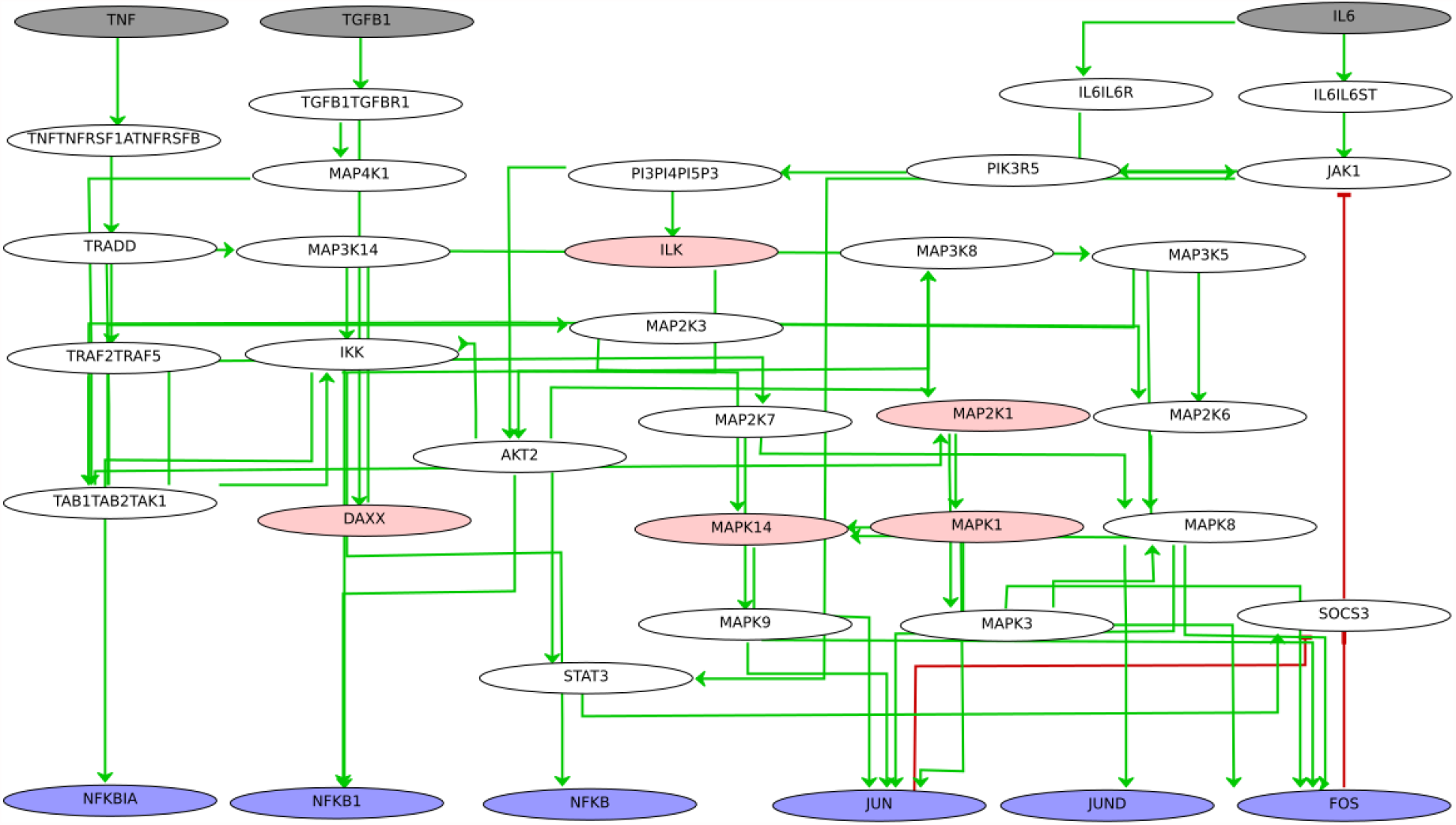
Boolean network comprising the three signalling cascades for TNF, TGFB1 and IL6. Boolean rules were inferred using the tool CaSQ (see Materials and Methods for a step by step model inference). Inputs are depicted in grey, TFs of interest are depicted in purple and intermediate nodes affected by the drug treatment or identified as mutation carriers are depicted in pink. Green arrows denote activation and red, blunt arrows denote inhibition.

The analysis shows that for the TFs identified as master regulators, the activation of IL6 or IL6 and TGFB1, is capable of positively regulating their expression, even in the presence of the anti-TNF treatment (TNF=0). Blocking the TNF cascade is able to completely shut down their expression only if combined with the blocking of IL6 and TGFB1 cascades. Regarding DAXX, it is actively expressed only when TGFB1 is activated and ILK is dependent on the activation of IL6. The MAPK molecules are dependent on the activation of IL6 and TGFB1 and do not seem to be impacted by the blocking of TNF.

Next, we created a reduced version of the Boolean modelling using the reduction function of the software GINsim in order to perform *in silico* experiments with combined perturbations. The reduced Boolean model comprised 23 nodes and for the analysis we created virtual KOs (knock outs) for a) MMP14 and MAPK1, b) DAXX, ILK and MAP2K1, and c) DAXX and NFKB1 to mimic the effects of the anti TNF treatment and also the mutation carriers, identified previously. For the simulations we set the initial conditions for the TNF to zero, and let the other inputs free, while setting the initial condition for all intermediate nodes to zero as well.

The results of the *in silico* experiments are shown in Tables 5, 6 and 7, for the molecules of interest and for each of the three conditions, respectively. For each set of conditions, the system was able to reach three steady states. For the first set of conditions, we observe that besides DAXX that is strictly TGFB1 dependent (Table 5, ss2 and ss3), all TFs can get activated with the presence of IL6 or IL6 and TGFB1, despite the TNF blockage and the downregulation of MAPK14 and MAPK1.

**Table 5.** Stable states of the Boolean network (MAPK1, MAPK14 KO, input TNF=0)

| Steady states | TNF | IL6 | TGFB1 | JUN | FOS | JUND | NFKBIA | DAXX | ILK | NFKB1 | MAP2K1 | MAPK1 | MAPK14 |
| --- | --- | --- | --- | --- | --- | --- | --- | --- | --- | --- | --- | --- | --- |
| ss1 | 0 | 0 | 0 | 0 | 0 | 0 | 0 | 0 | 0 | 0 | 0 | 0 | 0 |
| ss2 | 0 | 1 | 0 | 1 | 1 | 1 | 1 | 0 | 1 | 1 | 1 | 0 | 0 |
| ss3 | 0 | 1 | 1 | 1 | 1 | 1 | 1 | 1 | 1 | 1 | 1 | 0 | 0 |

**Table 6.** Stable states of the Boolean network (DAXX, ILK and MAP2K1 KO, input TNF=0)

| Steady states | TNF | IL6 | TGFB1 | JUN | FOS | JUND | NFKBIA | DAXX | ILK | NFKB1 | MAP2K1 | MAPK1 | MAPK14 |
| --- | --- | --- | --- | --- | --- | --- | --- | --- | --- | --- | --- | --- | --- |
| ss1 | 0 | 0 | 0 | 0 | 0 | 0 | 0 | 0 | 0 | 0 | 0 | 0 | 0 |
| ss2 | 0 | 1 | 0 | 0 | 0 | 0 | 1 | 0 | 0 | 1 | 0 | 0 | 0 |
| ss3 | 0 | 1 | 1 | 1 | 1 | 1 | 1 | 0 | 0 | 1 | 0 | 0 | 1 |

**Table 7.** Stable states of the Boolean network (DAXX and NFKB1 KO, input TNF=0)

| Steady states | TNF | IL6 | TGFB1 | JUN | FOS | JUND | NFKBIA | DAXX | ILK | NFKB1 | MAP2K1 | MAPK1 | MAPK14 |
| --- | --- | --- | --- | --- | --- | --- | --- | --- | --- | --- | --- | --- | --- |
| ss1 | 0 | 0 | 0 | 0 | 0 | 0 | 0 | 0 | 0 | 0 | 0 | 0 | 0 |
| ss2 | 0 | 1 | 0 | 1 | 1 | 1 | 1 | 0 | 1 | 0 | 1 | 1 | 1 |
| ss3 | 0 | 1 | 1 | 1 | 1 | 1 | 1 | 0 | 1 | 0 | 1 | 1 | 1 |

For the second set of conditions, we observe that when TNF is blocked and DAXX, ILK and MAP2K1 are downregulated, IL6 signal alone is not enough to activate the TFs JUN, FOS, JUND and the kinases MAPK14 and MAPK1 (Table 6, ss2). When both IL6 and TGFB1 signals are on, all TFs are activated, and the activity level of kinase MAPK14 restored (Table 6, ss3).

Lastly, we simulated the effects of the mutation carriers, as identified by DisGeNet, and the TNF blockage on the activity of the TFs and the kinases in our network. In Table 7 we observe that despite TNF blockage and DAXX and NFKB1 downregulation, all identified TFs and kinases are activated in the presence of IL6 or for IL6 and TGFB1 combined activity.

## Discussion

In the present work we make efforts to combine gene co-regulation with mechanistic signalling cascades that provide information about upstream regulation. We make use of the integrative RA network to analyze transcriptomic data regarding anti-TNF treatment and also map information about known disease-associated mutation carriers. Lastly, we make use of the tool CaSQ to add Boolean dynamics to a subnetwork of interest to mimic the effects of the anti-TNF treatment and estimate the impact of IL6 and TGFB1, as well as the impact of the downregulated genes, on the activation profile of the identified TFs.

The nineteen TFs identified as master regulators, have been implicated in RA and/ or autoimmunity as supported by the literature evidence. Six out of the nineteen TFs were also present in the RA map, a state-of-the-art mechanistic network for the disease that was built using manual curation. These six TFs, namely JUN, JUND, FOS, NFKBIA, ETS1 and TNFAIP3 were used as a functional overlap between the co-regulation and the signalling events, enabling us to obtain a network comprising upstream cascades, active TFs and target genes.

We used the integrative network as a template to analyse two independent datasets regarding anti-TNF treatment. We observed downregulation of some of the TFs previously identified as master regulators. To study the impact of the treatment in parallel with the activity of other signalling cascades, we extracted subgraphs from the integrative network. We selected the cascades of TNF, IL6 and TGFB1, up to the first affected TF to reduce complexity and focus on the upstream regulators.

We selected IL6, as it is one of the targets of the biologic treatment in rheumatoid arthritis [36,41–43] and TGF beta because it is an immunomodulatory cytokine highly expressed in RA patients, with a role that is yet to be determined [44–46]. We adjusted the map to model framework described in Aghamiri et al [29], to obtain an executable Boolean subnetwork to perform *in silico* analysis. As demonstrated from the real time simulations and the dose response analysis, both IL6 and TGFB1 cascades could affect the expression of the TFs and, as seen from the component sensitivity analysis, IL6 and TGFB1 could even counterbalance the downregulation of the studied TFs caused by the TNF blockage.

The steady state analysis confirmed the real time simulation results showing that for the TFs identified as master regulators, the activation of IL6 or IL6 and TGFB1 cascades is capable of positively regulating their expression. Blocking the TNF cascade is able to completely shut down the expression of these TFs only if combined with the blocking of IL6 and TGFB1 cascades. Towards this direction, dual targeted therapies have been proposed, either with the development of dual target agents, blocking simultaneously IL6 and TNF for example, [47] or by administering two biologics at the same time. However, the administration of combined biologics has been linked to increased adverse effects and it is currently under study to evaluate better dosage schemes [48,49].

Simulations with combined KOs mimicking the effect of anti-TNF treatment in combination with the downregulation of genes observed in the analyzed datasets, confirmed the dependency of the TFs activation state on the presence of inputs, and further highlighted specific conditions. For example, when TNF is blocked and DAXX, ILK and MAP2K1 are downregulated, IL6 signal alone is not enough to activate the TFs JUN, FOS, JUND and the kinases MAPK14 and MAPK1 (Table 6, ss2). However, when both IL6 and TGFB1 signals are on, all TFs are activated, and the activity level of kinase MAPK14 restored (Table 6, ss3). MAPK14 (p38a kinase) and MAPK1 (ERK2), are two proteins known to play a pivotal role in RA and are activated by a variety of signals, including cytokines such as TNF and IL6, but also TGF beta [50]. While p38 had been proposed as a potential target to reduce destruction of bone and cartilage, p38 inhibitors have given disappointing results in regards to therapeutic efficacy [51,52].

As seen in Table 5, suppression of MAPK14 (p38a) does not inhibit the activation of the identified as master regulators TFs, even in the presence of anti-TNF treatment, as other inputs, such as IL6 or TGF beta can counterbalance the effects. Regarding ERK inhibitors, limited data are available which may be due to a lack of efficient pharmacological inhibitors [53]. In older studies, FR180204, an ERK inhibitor, had demonstrated effectiveness against mouse collagen-induced arthritis [54], but there was no significant follow up. In our model, ERK2 inhibition (MAPK1) does not seem to significantly impact the activation of its downstream target TFs, JUN and FOS, as other regulators are also able to activate them.

While resistance to TNF therapy is a common event in the treatment of RA, the reasons behind its mechanisms are still unclear [55]. Currently there is no way of predicting which patient will respond or not to targeted therapy [56]. Executable, integrated networks can help towards personalized models, as mapping dysregulated genes could reveal potential impacted pathways shedding light onto the response to therapy. Dynamic analysis and *in silico* simulations can also inform about the outcome of combined perturbations, predicting the emergent behaviour of the system. Integrating multi-omic data is a key step in understanding pathogenetic mechanisms of multifactorial diseases, where one level of information does not suffice to explain the complex phenotypic traits. Patient-level analysis, by projecting patient-specific data onto a global network, and analysing the effects of patient-specific mutations and DEGs, in combination to treatment effects, could inform about the possibilities of success of a given therapy. Larger patient cohorts and more efficient computational techniques that would allow simulations in a greater scale could enhance the robustness and the predictive power of such models, helping to understand the response or non response to a given therapy at a patient level.

## Supplementary Materials

Supplementary Figures and Table are also available here for review purposes: https://docs.google.com/document/d/1pW7EuVMr4WOK4qyV2SDVOr8-pMle7Dbp9TK0EXNhs8o/edit

**Figure S1. Principal component analysis (PCA) in human blood cells with affected and unaffected Rheumatoid arthritis patients**.

The PCA shows 95 samples whose matrix expression data was downloaded from the GEO database (GSE117769). A variance stabilizing transformation was carried on matrix expression data.

**Figure S2. Global network for RA with associated DEG from responders/non responders RA patients data to anti-TNF treatment**.

Overlapping RA DEG from Adalimumab treatment and Etarnecept treatment data are shown in brown (11) and pink (4) respectively while non-overlapping genes/proteins are shown in grey (599).

**Figure S3. Global network for RA with associated DEG from untreated and treated RA patients data to anti-TNF treatment**.

Overlapping RA DEG are shown in green (101) while non-overlapping genes/proteins are shown in grey (513).

**S4 Fig. Environment sensitivity analysis. TFs could be upregulated in the absence of TNF activity, for a combination of activity ranges of the other two inputs (IL6 and TGFB1)**.

Upper part of each image corresponds to the correlation (positive above 0.0 and negative below it) of the sensitivity of the external components (shown in blue box) to optimize the activation of chosen internal components (TFs here). Lower part of the image corresponds to the box plots to show the range of activity percentage of external components (IL6 in blue, TGFB1 in orange and TNF as black line as it is chosen to be off) for the optimization of the chosen internal components activation. **(a)** optimize JUND activation. **(b)** optimize JUN activation. **(c)** optimize FOS activation. **(d)** optimize NFKB1 activation. **(e)** optimize NFKBIA activation.

**Table S1. Transcription factors identified from CoRegNet and their involvement in RA based on literature evidence**.

## Author Contributions

Conceptualization, A.N.; methodology, A.N. and M.E.; software, Q.M., D.D.M., V.S., M.E and A.N.; validation, A.N., Q.M. and V.S.; formal analysis, Q.M., D.D.M., M.E. and A.N.; investigation, Q.M., D.D.M. M.E. and A.N.; resources, V.C., E.P.T. and A.N.; data curation, Q.M., D.D.M and V.S.; writing—original draft preparation, Q.M., D.D.M., V.S. and A.N.; writing—review and editing, Q.M., D.D.M., V.S., V.C., M.E., E.P.T. and A.N.; visualization, Q.M. and V.S.; supervision, M.E. and A.N.; project administration, A.N.; funding acquisition, V.C., E.P.T and A.N. All authors have read and agreed to the published version of the manuscript.

## Data Availability Statement

Datasets (GSE117769, GSE129705, GSE138747 and DisGeNET variants) used for the analysis are publicly available. All data and code used to generate results, including networks inference, differential expression analysis, network visualisation is available on a GitLab repository at https://gitlab.com/genhotel/inference-of-a-global-integrative-network-for-rheumatoid-arthritis

The Shiny app login is: **genhotel** and the password is: **3886**

## Acknowledgements

We would like to thank Dr Smahane Chalabi, GenHotel, UEVE for her comments on the statistical treatment of the dataset used to infer the CoRegNet object.

