## Supplemental data for "Inference of an integrative, executable network for Rheumatoid Arthritis combining data-driven machine learning approaches and a state-of-the-art mechanistic disease map"

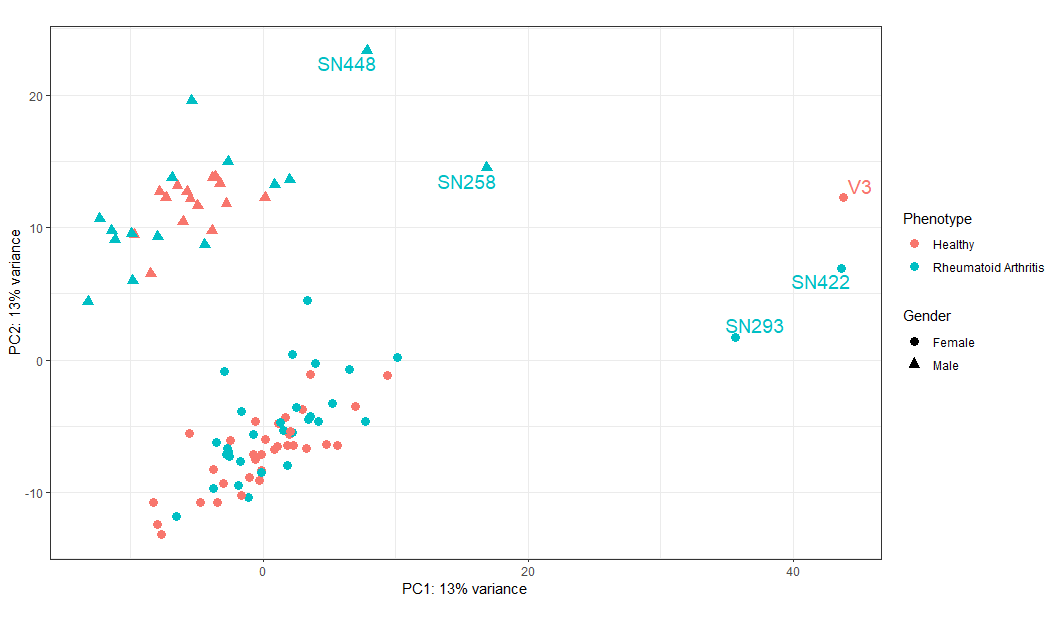

**S1 Fig. Principal component analysis (PCA) in human blood cells with affected and unaffected Rheumatoid arthritis patients.**

The PCA shows 95 samples whose matrix expression data was downloaded from the GEO database (GSE117769). A variance stabilizing transformation was carried on matrix expression data.

**S1 Table. Transcription factors identified from CoRegNet and their involvement in RA.**

| **No** | **Transcription factors** | **Role in RA** | **References** |
| --- | --- | --- | --- |
| **1** | **TNFAIP3** | NF-κB target gene, also involved in negative-feedback mechanism to block NF-κB activation through its ubiquitin-editing function in response to various inflammatory signaling, including TNF, IL-1β | PMID: 20822710  PMID: 22402800  PMID: 20852893  PMID: 26405544 |
| **2** | **IRF1** | IRF1 is critical for the TNF-driven interferon response in rheumatoid fibroblast-like synoviocytes | PMID: 31285419  PMID: 21834067  PMID: 32765497 |
| **3** | **ETS1** | Factor involved in the cytokine-mediated inflammatory and destructive cascade which is a characteristic of RA | PMID: 23101665  PMID: 11229456  PMID: 11976735 |
| **4** | **FOS** | Subunit of AP1 transcription factor which is involved in the transcriptional regulation of many pro inflammatory genes in RA | PMID: 19395871  PMID: 8660103  PMID: 9153554 |
| **5** | **NFKBIA** | Involved in different pathways and cellular processes such as TNFα signalling via NFκB | PMID: 30468518  PMID: 18454843 |
| **6** | **JUND** | Subunit of AP1 transcription factor which is involved in the transcriptional regulation of many pro inflammatory genes | PMID: 9764613  PMID: 17515956 |
| **7** | **HCLS1** | Dysregulated in RA synovial tissue | PMID: 12905466  PMID: 19563633 |
| **8** | **SPI1** | Essential for the expression of gliostatin/thymidine phosphorylase in RA which has angiogenic and arthritogenic activities | PMID: 22534375  PMID: 28192374 |
| **9** | **MXD1** | Expressed in RA peripheral blood cells, RA synovium | PMID: 22753658  PMID: 10568429 |
| **10** | **JUN** | Subunit of AP1 transcription factor which is involved in the transcriptional regulation of many pro inflammatory genes in RA | PMID: 18454843 |
| **11** | **NFKBIZ** | Involved in TNF and IL-17 mediated signaling | PMID: 32079724 |
| **12** | **TRIM22** | Expressed in RA peripheral blood | PMID: 24756903 |
| **13** | **FOSB** | Subunit of AP1 transcription factor which is involved in the transcriptional regulation of many pro inflammatory genes in RA | PMID: 29326694 |
| **14** | **DDX5** | DDX5 is required for the transcription of key Th17 genes involved in Th17-mediated autoimmune inflammation in RA | PMID: 29254845 |
| **15** | **BCL6** | Interleukin-29 regulates T follicular helper cells by repressing BCL6 in RA | PMID: 16508929  PMID: 28150777  PMID: 32468318 |
| **16** | **MNDA** | Citrullinated protein identified in RA synovial fluid; Interferon induced nuclear and cytoplasmic protein | PMID: 23044660  PMID: 15158620 |
| **17** | **EEF1A1** | Expressed in RA peripheral blood | PMID: 21444302 |
| **18** | **PTMA** | Regulated by c-Myc, an oncoprotein overexpressed in synovium of RA, and is associated with cell proliferation | PMID: 17372028 (mice) |
| **19** | **DAZAP2** | Expressed in RA peripheral blood mononuclear cells (PBMCs) | PMID:26352601 |

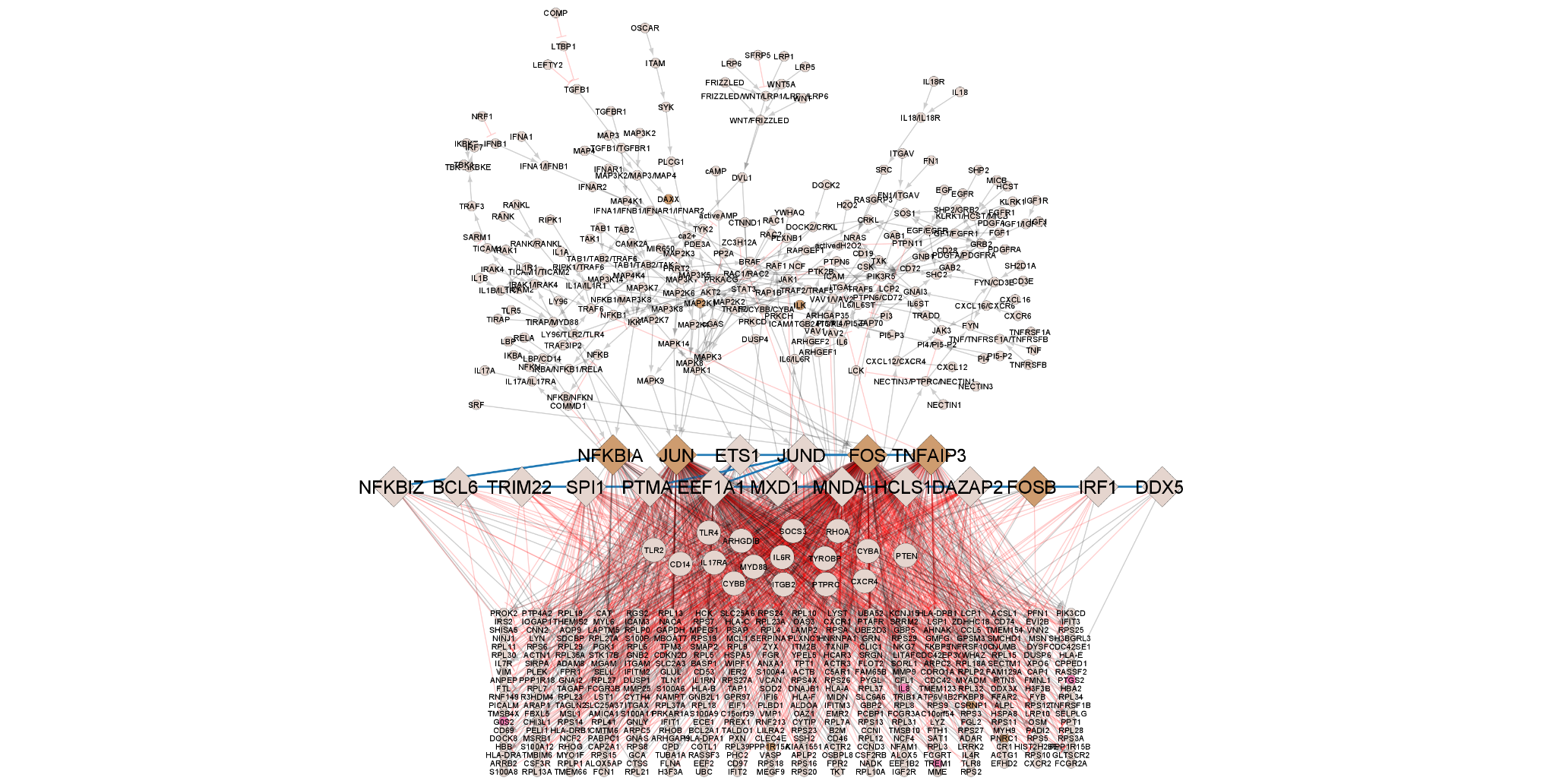

**S2 Fig. Global network for RA with associated DEG from responders/non responders RA patients data to anti-TNF treatment.** Overlapping RA DEG from Adalimumab treatment and Etarnecept treatment data are shown in brown (11) and pink (4) respectively while non-overlapping genes/proteins are shown in grey (599).

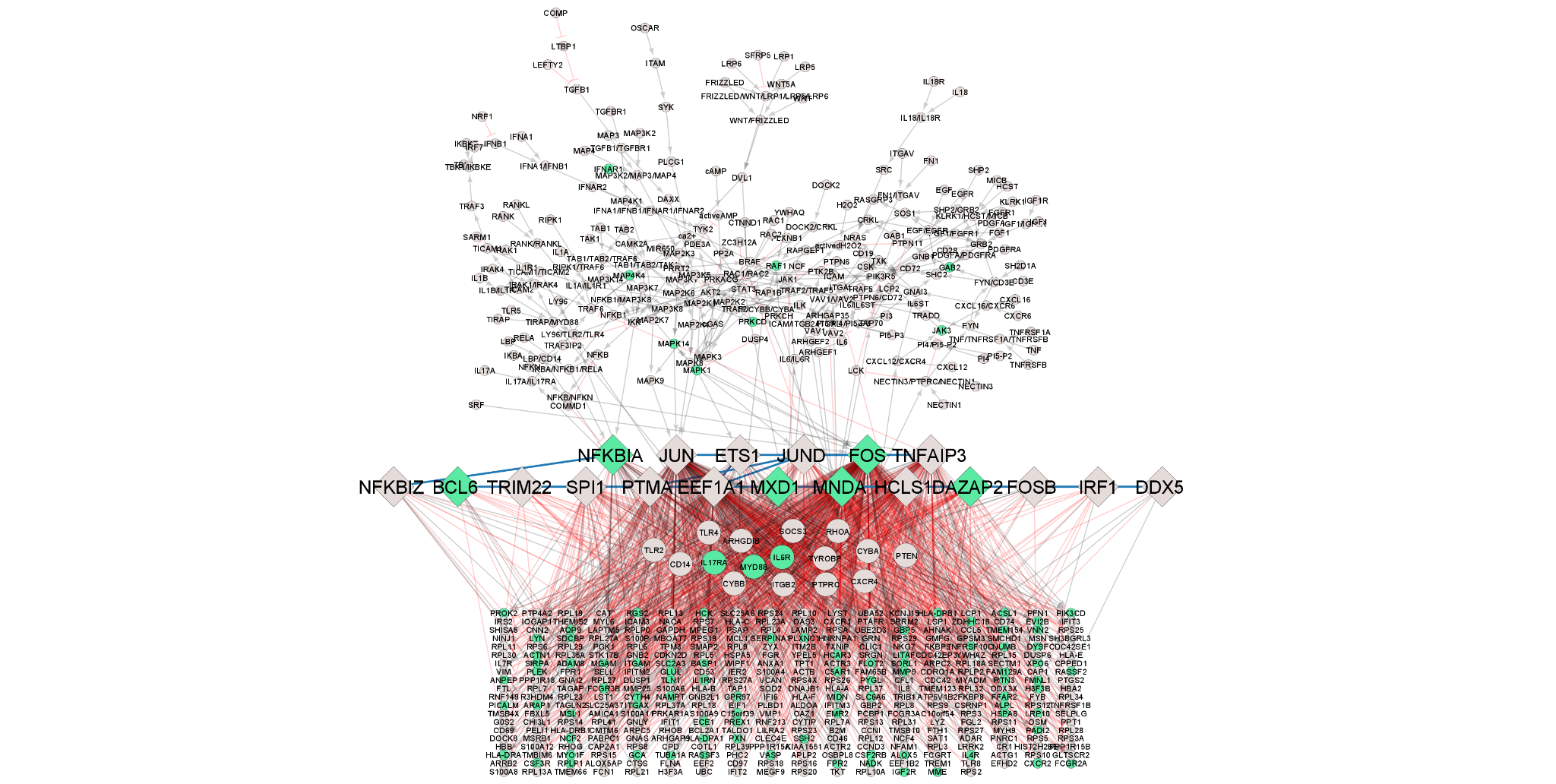

**S3 Fig.** **Global network for RA with associated DEG from untreated and treated RA patients data to anti-TNF treatment.** Overlapping RA DEG are shown in green (101) while non-overlapping genes/proteins are shown in grey (513).

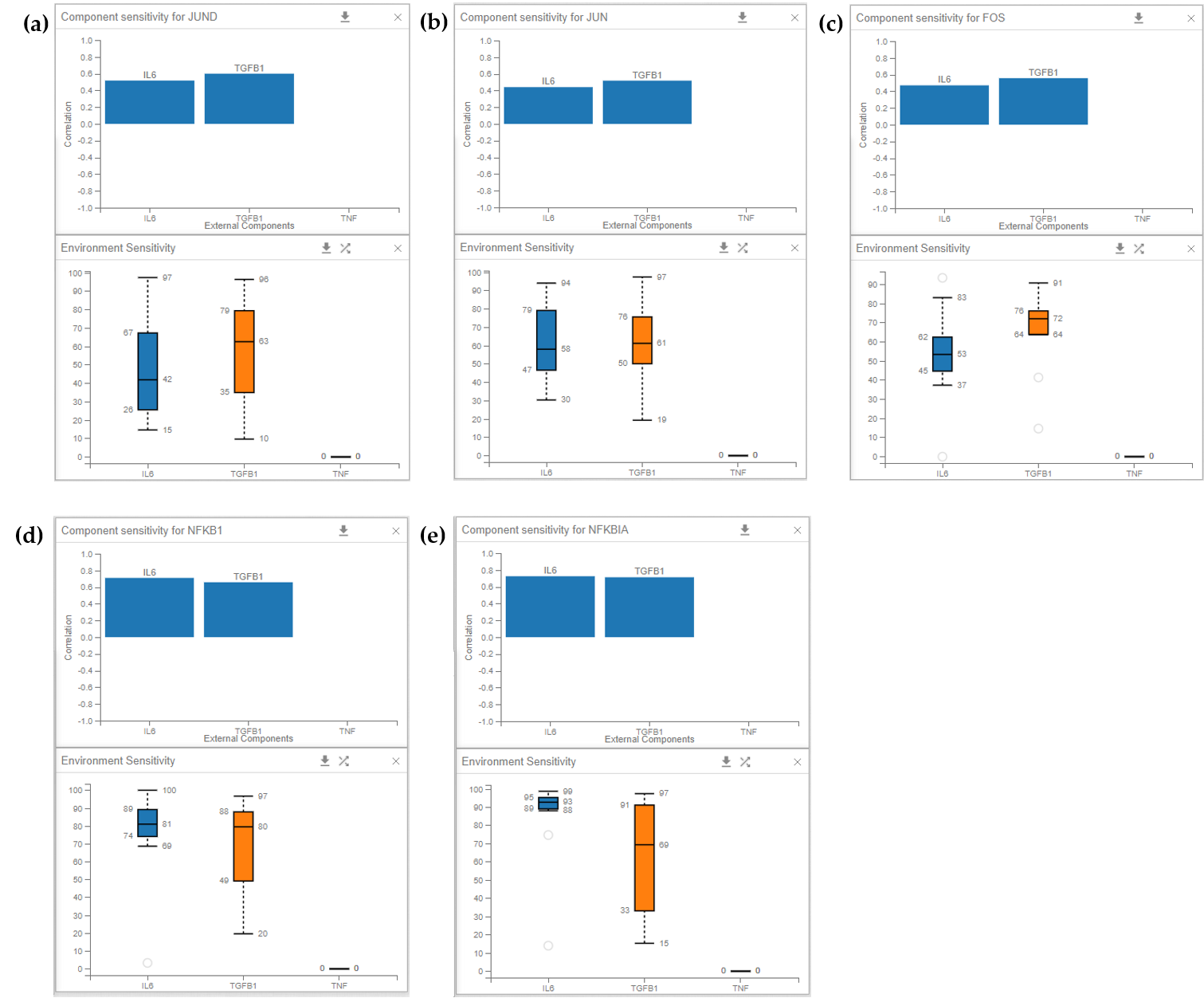

**S4 Fig.** **Environment sensitivity analysis. TFs could be upregulated in the absence of TNF activity, for a combination of activity ranges of the other two inputs (IL6 and TGFB1).** Upper part of each image corresponds to the correlation (positive above 0.0 and negative below it) of the sensitivity of the external components (shown in blue box) to optimize the activation of chosen internal components (TFs here). Lower part of the image corresponds to the box plots to show the range of activity percentage of external components (IL6 in blue, TGFB1 in orange and TNF as black line as it is chosen to be off) for the optimization of the chosen internal components activation. **(a)** optimize JUND activation. **(b)** optimize JUN activation. **(c)** optimize FOS activation. **(d)** optimize NFKB1 activation. **(e)** optimize NFKBIA activation.
